# Defining and Characterizing the Accessory Genome of the Virulent Vibriophage ICP1

**DOI:** 10.64898/2026.11.26.769439

**Authors:** Elizabeth A. Tan, Hugh B. Lawrence, Ariana C. Calderón-Zavala, David W. Lazinski, Andrew Camilli

## Abstract

Bacteriophages (phages) must evade host defense systems and redirect cellular resources to support viral replication. The virulent phage ICP1 is a persistent predator of *Vibrio cholerae* in both human hosts and aquatic environments, yet the genetic factors underlying its infection cycle remain incompletely defined. Here, we developed a transposon mutagenesis platform for ICP1 and applied transposon sequencing (Tn-seq) to identify genes required for infection across host strains and environmental conditions. Approximately 32% of ICP1’s 237 genes appeared essential, while 68% were dispensable, with a subset showing conditional essentiality. As a proof of principle, mutants disrupted in the anti-PLE (phage-inducible chromosomal-like element) counter-defense gene adi (gp99.1) were conditionally restricted in hosts carrying PLE2 when CRISPR-Cas-mediated targeting of PLE2 was abrogated but were otherwise neutral. We further identified phage genes involved in nucleotide metabolism whose contribution to fitness depended on nutrient availability. Under nutrient-limited conditions, mutants in the aerobic and anaerobic ribonucleotide reductase genes, *nrdAB* and *nrdDG*, were strongly depleted, consistent with roles in maintaining nucleotide pools under differing oxygen concentrations. Mutations in gp170, encoding a predicted dNMP kinase, and gp193 (*td*), encoding a predicted thymidylate synthase, also resulted in fitness defects. Together, these results help establish Tn-seq as a powerful approach for defining ICP1 gene requirements while highlighting the need to validate subtle conditional fitness effects.

**IMPORTANCE:** *V. cholerae* inhabits both aquatic reservoirs and the human host, exposing its phages to diverse nutrient environments and host physiological states that laboratory studies rarely examine. Tn-seq has recently been applied to study the gene requirements of virulent phages. In this study, we develop a novel Tn-seq method to study the virulent vibriophage ICP1 in distinct infection contexts, including during the stationary growth phase and under estuary-like conditions that better reflect the ecological settings of its host, *V. cholerae*. This approach revealed that genes involved in nucleotide metabolism can contribute to phage fitness under nutrient limitation. These findings demonstrate how genetic approaches can reveal conditionally important genes and provide a framework for studying viral fitness in the diverse environments occupied by bacterial hosts.

## INTRODUCTION

Phages are diverse and abundant viral predators of bacteria. Phages must overcome a myriad of host-encoded defense systems that vary by strain while redirecting cellular resources toward progeny production. This evolutionary arms race has generated a wide array of molecular strategies on both sides (1). However, the functions of most phage genes remain uncharacterized, in part due to a paucity of functional genomic tools capable of genome-wide screens.

Essential genes required for replication and virion assembly make up the conserved core genome, while accessory genes that vary between phage isolates make up the pangenome. It is hypothesized that accessory genes expand host range and support replication in diverse environments (2, 3). For example, phage infection often occurs under nutrient-limited conditions or in the presence of host defense systems, both of which can reduce replication efficiency (4, 5). To overcome these constraints, phages encode conditionally important genes that support infection in diverse environmental and host contexts. Identifying these genes requires approaches that assess phage fitness across varying host genetic backgrounds and growth conditions.

Genome-wide approaches such as Tn-seq have proven powerful for defining gene fitness contributions in bacteria. This method relies on sequencing transposon–genome junctions within a saturated mutant library to quantify the relative abundance of insertion mutants before and after selection. Changes in mutant frequency directly measure gene contribution to fitness, while essential genes are readily identified by their lack of transposon insertions (26). Adapting this approach to virulent phages is technically challenging because generating a transposon library is difficult: phage infection cycles are rapid, phages suppress host transcription, and antibiotic selection cannot select mutants. As a result, genome-wide functional studies in virulent phages have only recently become feasible (6–11).

The pathogen *V. cholerae* has a permanent aquatic reservoir and infects the human small intestine, where it causes cholera, a severe diarrheal disease endemic to regions such as South Asia and parts of Africa (12–14). Researchers frequently isolate the virulent vibriophage ICP1 from cholera patient stool samples and the environment in cholera-endemic countries (15, 16). ICP1 is unusual in that, unlike other virulent vibriophages, it can replicate in *V. cholerae* under both nutrient-rich and nutrient-poor conditions, including in estuarine environments and in stationary-phase cells (17). ICP1’s ∼126 kb genome contains conserved operons encoding essential replication and structural functions alongside a large accessory gene set largely of unknown function (18).

*V. cholerae* encodes a wide array of phage defense systems (19). ICP1, in turn, encodes counter-defense mechanisms to overcome these barriers. Many *V. cholerae* strains harbor a PLE that blocks ICP1 replication and redirects viral proteins for its own spread (20, 21). In response, ICP1 has evolved multiple mechanisms to counter PLEs, including a type I-F CRISPR-Cas system, the origin-directed nuclease Odn, and the PLE2-specific inhibitor Adi (22, 23). Additional ICP1 accessory genes counter other phage defense systems, such as the BREX inhibitor OrbA, further highlighting the diversity of phage counter-defense strategies (24). These interactions underscore the importance of phage genes that are required only in specific host or environmental contexts.

Here, we systematically identified ICP1’s essential core genes and conditionally important genes using Tn-seq. We generated a near-saturated transposon mutant library and quantified mutant fitness across multiple host strains and growth conditions. This approach enables genome-wide identification of genes required for phage replication across diverse infection contexts, providing new insight into phage adaptation to environmental and host-imposed constraints.

## MATERIALS AND METHODS

### Strains and growth conditions

Strains, plasmids, and phages used in this study are listed in **Tables 1-3**, respectively. Unless otherwise noted, *V. cholerae* was grown in lysogeny broth (LB) with aeration at 37 °C. Estuary-like conditions were modeled using Instant Ocean (IO; 0.7% w/v; Spectrum Brands Pet, LLC, Blacksburg, VA, USA) supplemented with ground shrimp chitin (1% w/v) and incubated at 30 °C with shaking. Antibiotics were used at standard laboratory concentrations unless noted otherwise. High titer phage stocks were generated by infecting *V. cholerae* at a multiplicity of infection (MOI) of 0.01 in LB and incubating for 4 h at 30 °C with aeration. Lysates were clarified by centrifugation at 12,000 × g for 10 min at 4 °C, and phages were precipitated overnight with 10% polyethylene glycol (MW 8000; PEG). Precipitated phages were pelleted by centrifugation at 10,000 × g for 20 min; the pellet was resuspended in IO and stored at 4 °C. Site-specific mutations were made in the *V. cholerae* genome using allelic exchange following natural transformation as described (42). We subjected all mutant strains to whole-genome sequencing to verify the mutations and confirm the absence of spontaneous mutations.

**Table 1.**
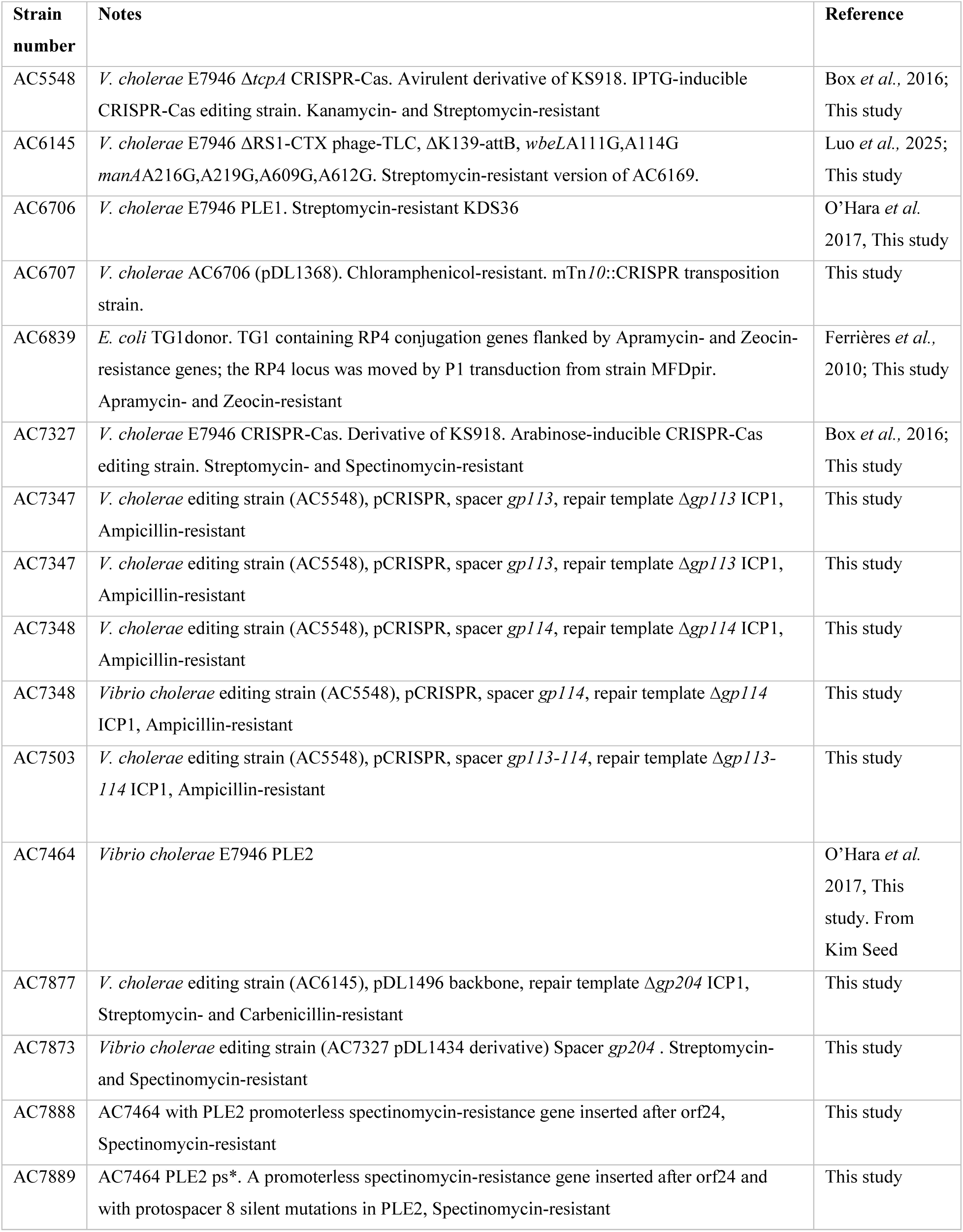
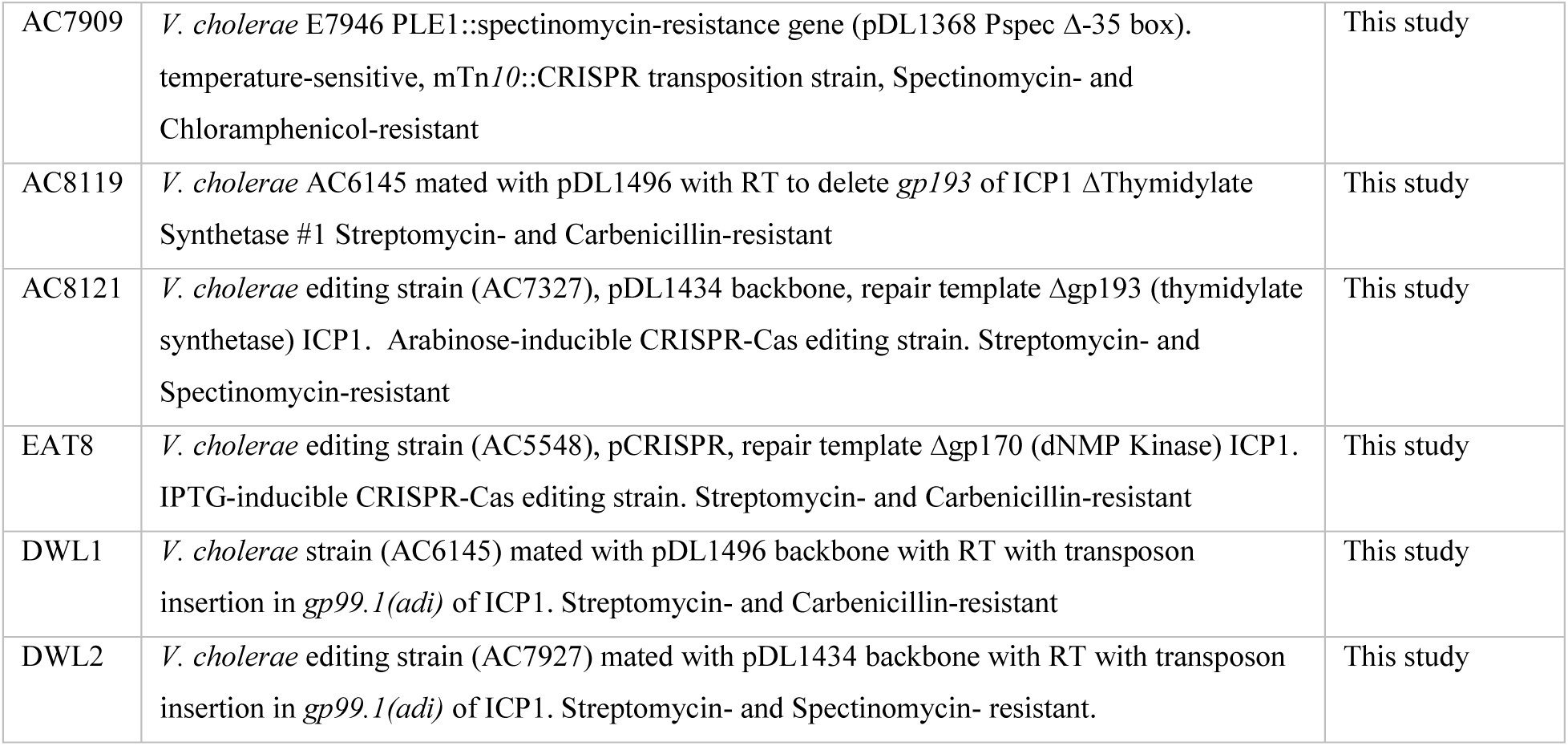
Strains used in this study.

**Table 2.**
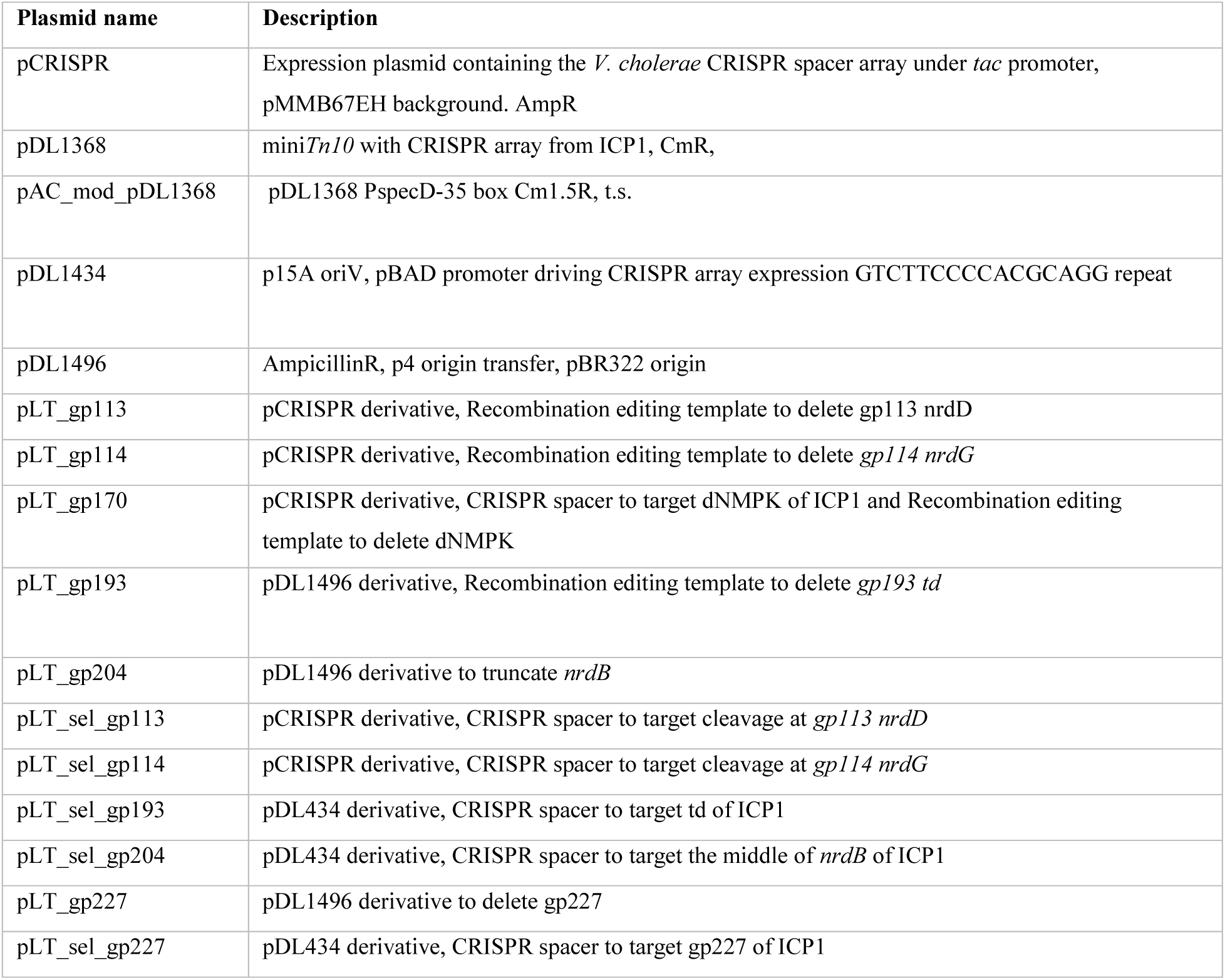
Plasmids used in this study.

**Table 3.** ICP1 isolates and mutants used in this study.

| **ICP1 isolates used** | **Description** | **Reference** |
|---|---|---|
| ICP1_2011_A | wild-type | clinical isolate, (15) |
| ΔS2-9 | deleted spacers in the CRISPR array, Cas1 mutant to prevent spacer acquisition | made by Kim Seed<br>(21) |
| Δ*nrdD* | Deletion mutant of anaerobic ribonucleotide reductase alpha subunit deleted, gp113 | this study |
| Δ*nrdG* | Deletion mutant of anaerobic ribonucleotide reductase beta subunit, gp114 | this study |
| Δ*nrdB* | Truncation mutant of *nrdB, gp204* | this study |
| adi:: Tn | cloned transposon mutant of *gp99.1* with mTn*10* with mini-CRISPR array, made in the ΔS2-9 background | this study (made by DWL) |
| Δ*dNMPK* | Deletion mutant of dNMP Kinase deleted, gp170 | this study |
| Δgp227 | Deletion mutant of Small conserved hypothetical gene of ICP1, gp227 |  |
| Δ*td* | Deletion mutant of *td* which encodes thymidylate synthetase, gp193 | this study |
| Δ*nrdDG* | Deletion mutant of anaerobic ribonucleotide reductase system deleted, gp113-114 | this study |

### Construction of plasmids

We constructed a high-temperature-inducible mTn*10* transposition plasmid as a derivative of pDL1093 (25). We made a minimal mTn*10* containing a mini-CRISPR array with two anti-PLE1 and PLE2 spacers and one anti-PLE5 spacer, driven by a constitutive promoter (TTGAACATTTTAATGGTAATGTGATAAAATAGGTAC). The resulting plasmid, pDL1368, was transformed into the PLE1-containing *V. cholerae* strain AC6706 by natural transformation with selection at 30 °C. Transcription of the transposase is repressed below 32 °C and induced maximally at 39 °C. A derivative of pDL1368, designated pAC_mod_pDL1368, was constructed where the −35 box in the promoter driving the mini-CRISPR array was deleted. To aid in making genome edits in ICP1, derivatives of pCRISPR with targeting spacers were constructed as described (27). We constructed an alternative CRISPR targeting plasmid, pDL1434, using Splicing-by-overlap-extension (SOE) PCR. It contains a p15A origin of replication, RP4 origin of transfer (oriT), kanamycin resistance gene, *araC*, and an arabinose-inducible promoter (P_BAD_) driving expression of a minimal CRISPR array (GenBank accession pending). Targeting spacers were incorporated into pDL1434 by PCR using 5’ phosphorylated primers with 5’ extensions complementary to the parental plasmid’s CRISPR repeats, followed by circularization by blunt-end ligation. To make site-specific mutations in ICP1, we constructed an allelic exchange plasmid, pDL1496, by replacing the *lacZ* gene and polylinker of pUC19 with oriT and an ampicillin resistance gene (GenBank accession pending). We used SOE-PCR with 5′-phosphorylated primers to generate an intermediate allelic exchange fragment with terminal sequences complementary to pDL1496. The pDL1496 vector was amplified by PCR using 5′-phosphorylated primers. The plasmid and allelic exchange fragment were assembled in a secondary SOE-PCR step to yield a linear plasmid derivative, which was circularized via blunt-end self-ligation. Ligated DNA constructs were transformed into chemically competent *Escherichia coli* TG1 donor cells (a TG1 derivative encoding RP4 conjugation machinery), and subsequently transferred into *V. cholerae* by mating on LB agar plates at 37 °C for 1 hour.

### Phage plaque assays

For spot assays, overnight cultures of *V. cholerae* were mixed with molten 0.5% LB soft agar cooled to 45 °C and poured onto LB agar plates to generate soft agar lawns. After solidification, 10 µL of ten-fold serial phage dilutions were spotted onto the surface, allowed to dribble across the face of the plate, and allowed to dry. Plates were incubated overnight at 37 °C before imaging and enumeration of plaque-forming units (PFUs). For embedded plaque assays, mid-exponential growth phase cultures of *V. cholerae* were infected with serial phage dilutions and allowed to adsorb for 10 minutes at room temperature. The infected cells were mixed with molten 0.5% LB soft agar and poured on LB agar plates. After solidification, plates were incubated overnight at 37 °C before imaging and PFU enumeration.

### Construction of the ICP1 transposon insertion libraries

We performed transposon mutagenesis *in vivo*, i.e., during ICP1 infection of an inducible transposition host strain. *V. cholerae* harboring PLE1 and carrying the temperature-sensitive plasmid pDL1368 (strain AC6707) was grown in LB with chloramphenicol (1.5 µg/mL) at 31 °C to early stationary phase. Cultures were infected with ICP1 ΔS2–9 (deleted for CRISPR spacers 2-9, several of which target PLE1 and PLE2) at an MOI of 0.1 and distributed into 384-well plates. Plates were shifted to 39 °C for 5 h to induce transposition and allow phage replication and host cell lysis. Progeny phage were transferred to 96-well plates containing early exponential-phase PLE1 *V. cholerae* (AC6706) using a 96-pin replicator and incubated overnight at 37 °C. Approximately 95% of wells produced confluent plaques when spotted on AC6706 lawns. Lysates from all 96-well plates were pooled and clarified by centrifugation. Phages in the supernatant were precipitated with PEG, pelleted, resuspended in IO, and stored at 4 °C. A separate library was generated using the −35 box deletion transposition plasmid, pAC_mod_pDL1368, harbored in AC7909.

### Passaging of the phage transposon library and transposon-sequencing

The library was passaged on AC6706 at MOI = 0.01 in LB to either early exponential-phase or stationary-phase, or in estuary-like conditions. For the estuary-like condition, stationary-phase cells were pelleted, washed, and resuspended in IO, adapted overnight at 30 °C with aeration, then grown for 5 h in IO + 1% shrimp chitin flakes (Sigma-Aldrich, St. Louis, MO, USA) before infection. Phage lysates were collected after 8 h for the LB cultures or after 5 h for the IO plus chitin cultures. Depending on expansion of the phage population, lysates were harvested after one or two passages. Phage DNA was extracted from input and output populations using the Wizard DNA Purification Kit (Promega Corporation, Madison, WI, USA). DNA samples were used for Tn-seq as described (26).

### Phage gene deletion mutant and transposon insertion construction

We generated phage harboring in-frame gene deletions or an mTn*10* disruption of the *adi* gene (gp99.1) using a variation of the previously described CRISPR-counter-selection method (27). Briefly, we used ICP1 to infect *V. cholerae* containing either a pCRISPR or pDL1496 derivative with the desired mutation construct, allowing allelic exchange via double-crossover homologous recombination. We selected recombinant phages by plaque assay on *V. cholerae* CRISPR-targeting strains (AC5548 or AC7327) harboring pCRISPR or pDL1434 derivatives encoding spacers complementary to the wild-type allele. We purified selected plaques at least twice, screened them by PCR, and verified them by whole-genome sequencing.

### Phage competition

Competitions were performed between ICP1 ΔS2-9 and an *adi* transposon insertion mutant. Three independent input phage mixtures were generated containing either 1 or 0.1% *adi*::Tn. Each input was used to infect either a permissive PLE2 host or the PLE2 protospacer mutant strain for a single passage. For this, *V. cholerae* was grown at 37 °C with aeration to an OD_600_ of 0.1 in LB before adding phage at an MOI of 0.01. The culture was incubated for 2 hours at 37 °C until lysis was observed. Phage lysates were serially diluted and spotted onto lawns of either a PLE1 host, conditionally permissive for *adi*::Tn, or a host permissive for both ΔS2-9 and *adi*::Tn mutants. We compared the titers of these phage populations with the Tn-seq results.

## RESULTS

### Construction of transposon mutant library in ICP1

To systematically interrogate gene function in ICP1, we developed an *in vivo* transposon mutagenesis method and paired it with Tn-seq. We selected mTn*10* insertion mutants carrying a conditionally essential anti-PLE CRISPR array, allowing transposon mutants to replicate on a PLE host. Initial attempts to generate the library *en masse* resulted in libraries highly skewed by jackpot transposition events. To overcome this, we induced transposition separately in 1,152 wells across three 384-well plates. Because the mini-CRISPR array on the transposition plasmid is constitutively expressed, we did not select for transposon insertions during this step. We selected for phages with transposon insertions by passaging the phage from each of the 1,152 wells on PLE1 strain AC6706, which lacked the transposition plasmid. We performed this selection in 96-well plates. To construct a near-saturating library of transposon mutants, lysates from five biological replicates were pooled following selection.

### Genome-wide essentiality mapping of ICP1

Phage DNA was extracted from ∼10^9^ PFU of the library. We prepared a sequencing library by amplifying the transposon–genome junctions, then performed massively parallel sequencing on the Illumina platform. We obtained 103,374,959 reads, of which approximately 70% aligned to the ICP1 ΔS2–9 reference genome. From these data, we identified 2,189 unique insertion mutants supported by ≥5 reads (**Figs. 1 and S1**), which is one insertion every 58 bp on average.

**Figure 1.**
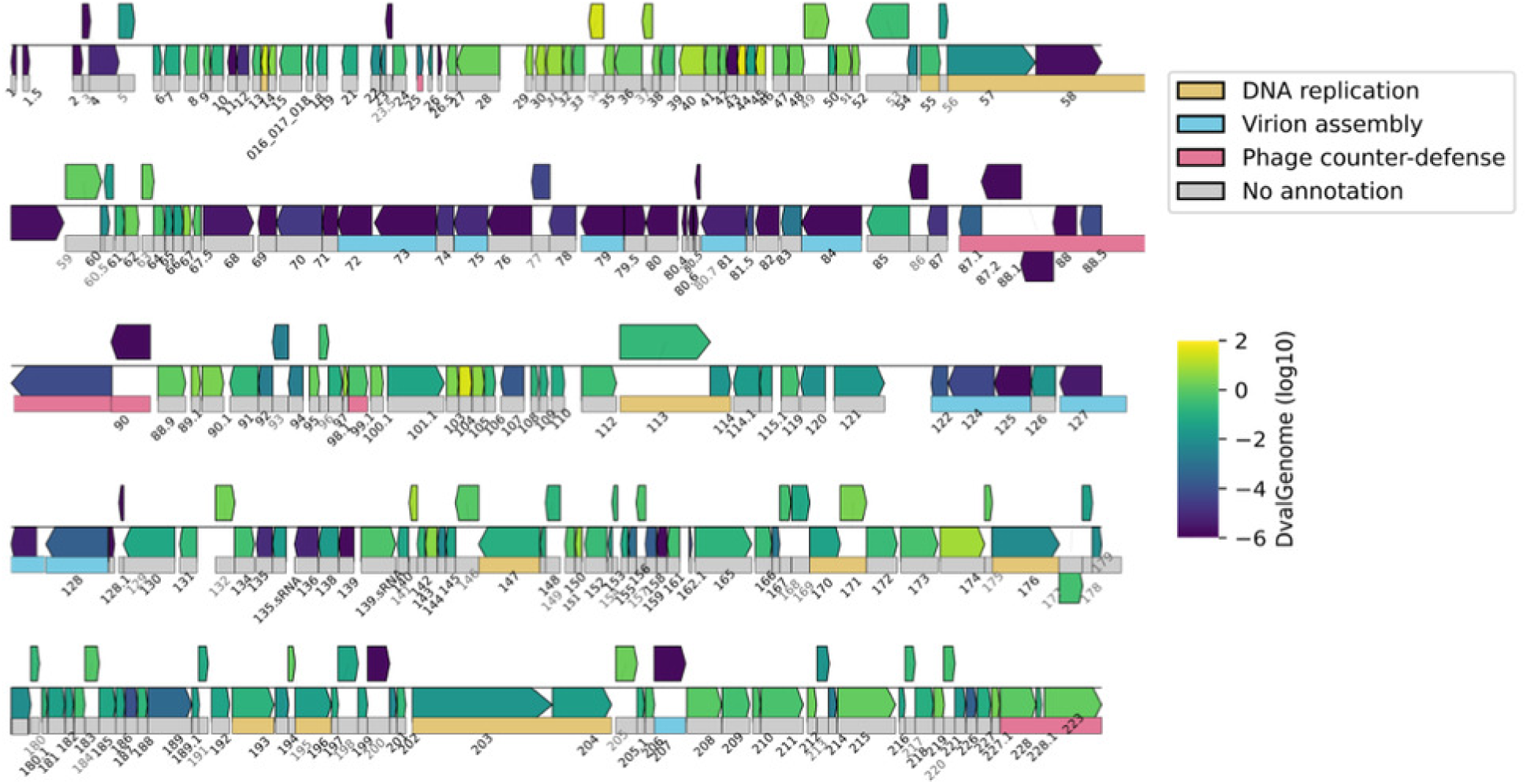
Phage gene fitness categorization. The ICP1 genes are represented as polygons pointing in the direction in which they are transcribed and numbered. Overlapping genes are vertically offset. The rectangles below each gene indicate the functional class according to the upper key. Gray rectangles indicate hypothetical genes with no known function. The color of each gene polygon indicates the fitness value or DvalGenome(log10) according to the lower key. Abundant mutants are lighter green, and rare mutants/putatively essential genes are dark purple.

We categorized ICP1 genes into four groups: advantageous, neutral, disadvantageous, and essential, based on their DvalGenome value, which is defined as the ratio of observed to expected read counts for each gene, where expected reads = total read count × (gene length/genome length). Genes with low DvalGenome values exhibit fitness defects when disrupted, whereas high DvalGenome values indicate a fitness advantage. Most genes had DvalGenome values <1, so we classified these mutations as disadvantageous or putatively essential (hereafter referred to as essential).

Across the 237 annotated ICP1 genes, we classified 76 (32%) as essential (**Fig. 1**). Although most of these lacked insertions, some contained one or a few insertions with extremely low read counts, consistent with a severe fitness defect. Given the limited number of ICP1 generations during library construction and selection, plus the potential for some mutants to be complemented *in trans* during coinfection in later rounds of infection when the MOI exceeds 1, we defined putatively essential genes as those having a DvalGenome value <0.02. The proportion of dispensable genes is consistent with that of other large genome phages such as Phikzvirus phiKZ and Tequatrovirus T4 (70% and 71.5% dispensable, respectively) (7, 8). As expected, genes encoding core replication machinery and virion structural proteins were essential. Because library selection required ICP1 CRISPR-Cas activity to restrict PLE, we also expected *cas* genes to be conditionally essential, which was the result for all six *cas* genes (**Fig. 1**). Several operons contained both essential and dispensable genes (*e.g.*, gp40-gp46 and gp134-139), suggesting functional modularity within transcriptional units. The library additionally revealed essential genes of unknown function, such as gp43. In contrast, adjacent genes in this region were dispensable, showing that this wasn’t simply a cold spot for mTn*10* insertion (**Fig. 1**).

We observed a strong directional bias in transposon insertion orientation, with most insertions aligning the constitutive CRISPR array promoter within mTn*10* with transcription of the disrupted gene or operon (*e.g.*, gp40-gp46; **Fig. S2**). This constitutive promoter is expected to generate outward-reading transcription into flanking phage genes. This directional bias likely results from a combination of fitness costs imposed by antisense transcription of genes and polarity on downstream genes. Both of these fitness costs are expected to be relieved when the transcriptional orientation of mTn*10* is the same as the disrupted gene or operon.

### Identifying conditionally important accessory genes of ICP1

Since *V. cholerae* strains often differ in their anti-phage defense repertoire and can frequently occupy nutrient-poor aquatic habitats, ICP1 may encode conditionally important genes that maximize replication efficiency in different hosts and under nutrient-poor conditions. To validate whether we could identify conditionally important phage counter-defense genes by Tn-seq, we designed a screen to measure the importance of *adi* (gp99.1), a known inhibitor of PLE2 (23). Adi functions redundantly with ICP1’s CRISPR-Cas system to inhibit PLE2. Therefore, we hypothesized that transposon disruption of *adi* would be fully restricted on a PLE2 host in the absence of the phage CRISPR-Cas activity. To test this, we abrogated CRISPR-Cas targeting by constructing host strain PLE2 ps*, which contains eight silent mutations in the PLE2 protospacer sequence. We passaged the ICP1 transposon library through *V. cholerae* strains harboring PLE1, PLE2, or PLE2 ps*. As expected, transposon insertions in *adi* or in the upstream gene gp098.1, which appears to be in an operon with *adi*, were depleted ∼8-fold and ∼4-fold after passage through the PLE2 ps* strain, respectively (**Fig. 2A**). In contrast, *adi* was neutral during infection of the PLE2 wild-type host (**Fig. 2B**). These results confirmed that Tn-seq can be used to identify conditionally important genes. Other mutants, including insertions in hypothetical genes gp101.1 and gp227, were also depleted following passage through PLE2 ps*. However, insertions in both genes were also depleted after infecting the PLE1 host during exponential growth, suggesting their contributions to phage fitness are not specific to the PLE2 ps* host. To further validate the Tn-seq results, we generated an in-frame deletion in gp227 and showed that the Δgp227 mutant formed smaller plaques than wild-type ICP1 (**Fig. S3**).

**Figure 2.**
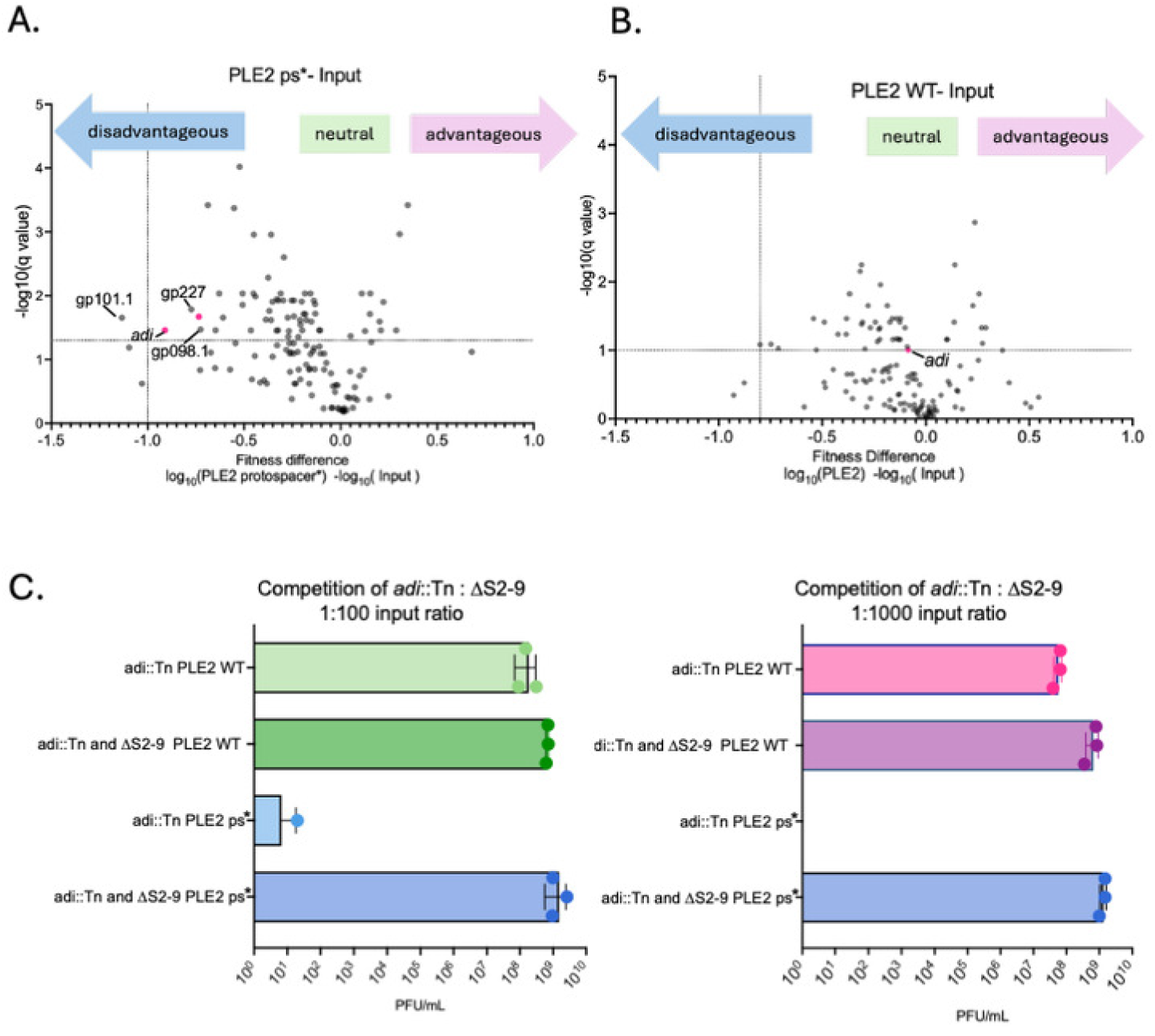
Tn-seq identifies a known conditionally essential gene of ICP1 for restricting PLE2. The ICP1 transposon mutant library was infected at an MOI = 0.01 on either a PLE2 variant strain carrying a mutant protospacer (PLE2 ps*) **(A)** or a wild-type PLE2 **(B)** in LB during exponential growth. Five biological replicates of phage lysates were collected and processed for Tn-seq. Volcano plots show the difference in log₁₀(DvalGenome) values between output and input populations (x-axis) and the – log₁₀(q value) (y-axis) after multiple hypothesis testing correction using the two-stage step-up false discovery rate (FDR) procedure (Benjamini, Krieger, and Yekutieli) with a desired FDR (Q) of 5%. Genes with higher y values are more confidently associated with differential fitness. The *adi* (gp99.1) gene and other genes (gp098, gp101.1, gp227) exhibited significant depletion in the PLE2 ps* condition, indicating conditional importance for antagonizing PLE2. **(C)** To validate the conditional requirement of *adi*, mixtures of *adi*::mTn10 and ΔS2-9 phages at 1:100 and 1:1000 ratios were prepared and infected in biological triplicates at MOI = 0.01 in either the WT PLE2 host or the PLE2 ps* (protospacer mutant). Phage lysates were titered individually for *adi*::Tn or for both phages, *adi*::Tn and ΔS2-9. The host is shown to the immediate left of each horizontal data column. The *adi*::Tn mutant is restricted on the PLE2 ps* host, confirming its conditional requirement.

As previously reported (23), ICP1 ΔCRISPR Δ*adi* cannot form plaques on a PLE2 host, so we next asked why the DvalGenome value of *adi* in the PLE2 ps* Tn-seq screen showed only a modest ∼8-fold reduction in fitness (**Fig. 2A**). One possibility is that, although *adi* mutants fail to replicate in PLE2 ps*, their DNA may remain largely intact. Consistent with this, a previous study showed that, after infection of a PLE2 host, wild-type ICP1 makes up ∼90% of mapped genomic reads while a ΔCRISPR Δ*adi* mutant makes up ∼75% even though it cannot replicate (23). To confirm that *adi* transposon mutants cannot produce infectious progeny in the absence of CRISPR-Cas targeting during competition, we infected PLE2 or PLE2 ps* hosts at an MOI of 0.01 with 1:100 and 1:1000 mixtures of ICP1 *adi*::Tn and ΔS2-9 parent strain. The ΔS2-9 mutant encodes Adi and therefore infects both PLE2 and PLE2 ps* hosts, whereas the *adi*::Tn mutant, whose CRISPR-Cas system can target PLE2, can infect PLE2 but not the PLE2 ps* host. Consequently, we expect no remaining *adi*::Tn phage to be present following passaging through the PLE2 ps* host. Indeed, we only observed two PFU from the 1:100 mixture in one of our three biological replicates, resulting in an extremely low mean PFU count, and zero PFU from the 1:1000 mixture (**Fig. 2C**). These results show that the *adi::*Tn mutant phage does not produce viable progeny but that the depletion observed by Tn-seq underestimates the fitness cost of losing *adi,* presumably because non-infectious genomes remain in the sequenced DNA.

### Identification of ICP1 genes important for infection under nutrient stress

*V. cholerae* is a native inhabitant of estuaries, where it relies primarily on chitin, a relatively poor carbon and nitrogen source for metabolism (28). The stationary phase sigma factor RpoS is essential for chitin utilization and growth in *V. cholerae* (29), linking nutrient stress to environmental survival. Notably, *V. cholerae* growth on chitin induces natural competence, which can promote horizontal gene transfer (30). Interestingly, ICP1 can infect non-growing (LB, stationary-phase) and nutrient-stressed *V. cholerae* (from estuary), albeit with low efficiency (17). We hypothesized that ICP1 carries conditionally important genes used to redirect host resources and scavenge metabolites, particularly nucleotide pools, to support viral replication under these conditions. To identify ICP1 genes important for infection of nutrient-stressed or stationary-phase cells, we passaged the ICP1 transposon library through *V. cholerae* grown under three conditions: exponential or stationary phase in LB at 37 °C, and estuary-like medium consisting of 0.7% Instant Ocean (IO) supplemented with 1% shrimp chitin flakes (IOC) at 30 °C. We collected phage lysates after one passage on exponential-phase *V. cholerae*, or two passages for the remaining two conditions to compensate for reduced phage multiplication. Tn-seq fitness differences were assessed using log₁₀(DvalGenome) values and visualized as volcano plots comparing each output condition to the input library.

Two ICP1 nucleotide metabolism genes, gp193 (*td*, thymidylate synthetase) and gp204 (*nrdB*, aerobic ribonucleotide reductase [RNR] beta subunit), were disadvantageous during infection of exponential-phase cells, which suggests these genes are generally important for the phage infection cycle (**Fig. 3A**). These and three additional nucleotide metabolism genes, gp170 (dNMP kinase) and gp113-gp114 (*nrdDG*, anaerobic RNR alpha and beta-subunit), emerged as strong hits under nutrient stress conditions (**Fig. 3B and C**). Although classified as non-essential, insertions in these genes showed DvalGenome values reduced by up to two orders of magnitude. To validate these findings, we generated deletion mutants Δ*td*, ΔdNMPK, Δ*nrdDG*, Δ*nrdB*, and Δ*nrdDG/B* in the ΔS2-9 ICP1 strain background. Although one-step growth curves did not reveal significant growth defects in our mutants compared to the ΔS2-9 parental strain (**Fig. S4**), plaque assays showed that all four deletion mutants formed smaller plaques than the parent in both embedded plaque assays (**Fig. 4A, B, and D**) and spot plaque assays (**Fig. 4C**). Spot (surface) assays are oxygenated, while embedded assays generate an oxygen gradient, with oxygen highest at the surface and decreasing through the agar matrix. The Δ*nrdDG* (anaerobic RNR) mutant formed significantly smaller plaques than wild-type, and this difference was more pronounced in the embedded plaque assay (**Fig. 4D**). This suggests that both RNR systems contribute to phage fitness across oxygen gradients. We could not generate IOC soft agar lawns because chitin is insoluble; therefore, the relative importance of ICP1 genes in IOC versus LB exponential-phase or stationary phase is limited to the Tn-seq data. Together, these results indicate that RNRs enhance infection efficiency, particularly under nutrient-limited conditions.

**Figure 3.**
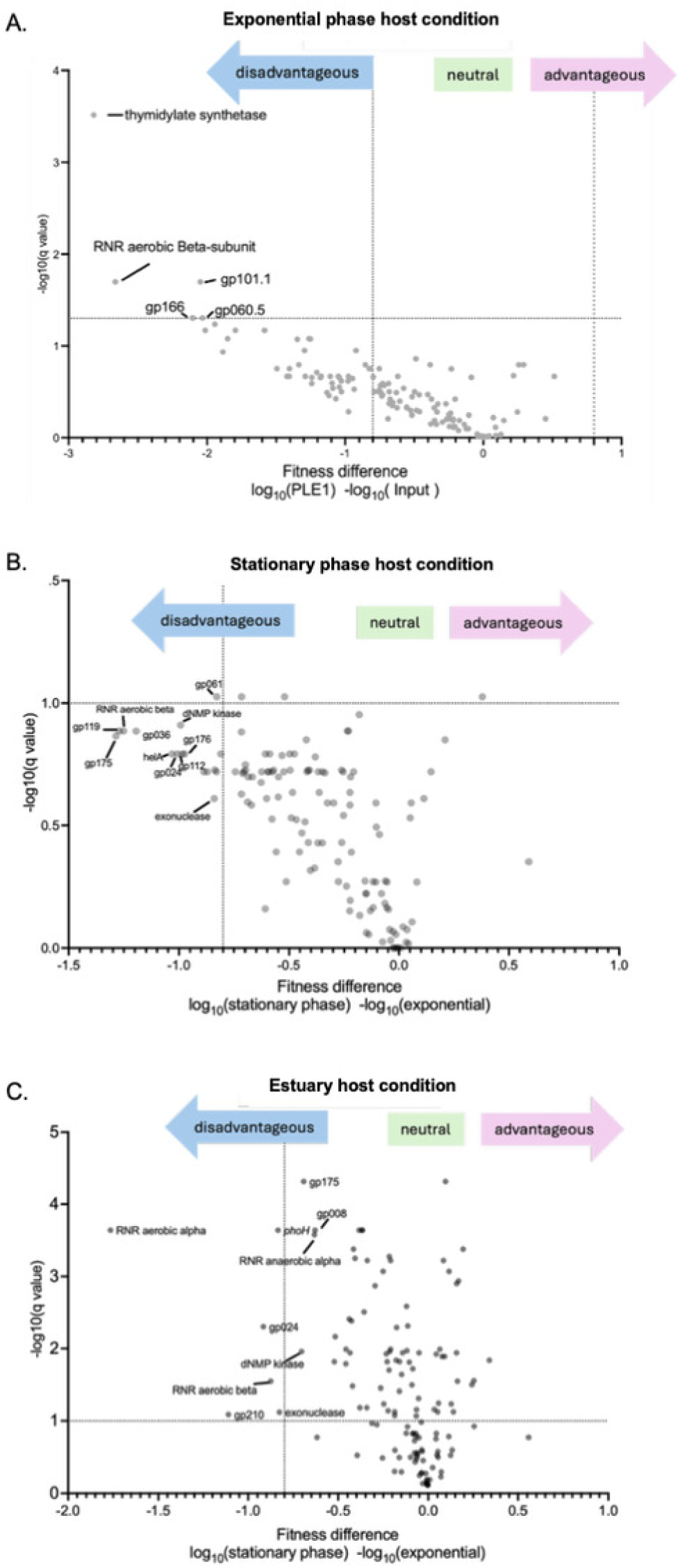
Nutrient-dependent fitness profiling reveals conditionally important nucleotide metabolism genes. The ICP1 transposon mutant library was used to infect a PLE1 variant strain under three conditions: **(A)** stationary phase in lysogeny broth (LB), **(B)** exponential phase in estuary-like Instant Ocean–chitin (IOC) medium, and **(C)** exponential phase LB. For conditions A and B, phage lysates were collected after two passages. For condition C, phage lysates were collected after one passage. Five independent biological replicates were processed for Tn-seq. Volcano plots show the difference in log₁₀(DvalGenome) values between output and input populations (x-axis) and the –log₁₀(q value) (y-axis) after multiple hypothesis testing correction using the two-stage step-up false discovery rate (FDR) procedure (Benjamini, Krieger, and Yekutieli) with a desired FDR (Q) of 5%. Genes exhibiting significant depletion included gp061, gp170, gp204, gp119, gp175, and gp036 under stationary-phase conditions (A); gp175, gp203, gp208 (*phoH*), gp170, and gp024 under estuary-like conditions (B); and gp193, gp204, gp101.1, and gp166 under exponential-phase conditions (C). These results identify genes associated with ICP1 fitness across distinct growth conditions, including genes involved in nucleotide metabolism and genes of unknown function.

**Fig. 4.**
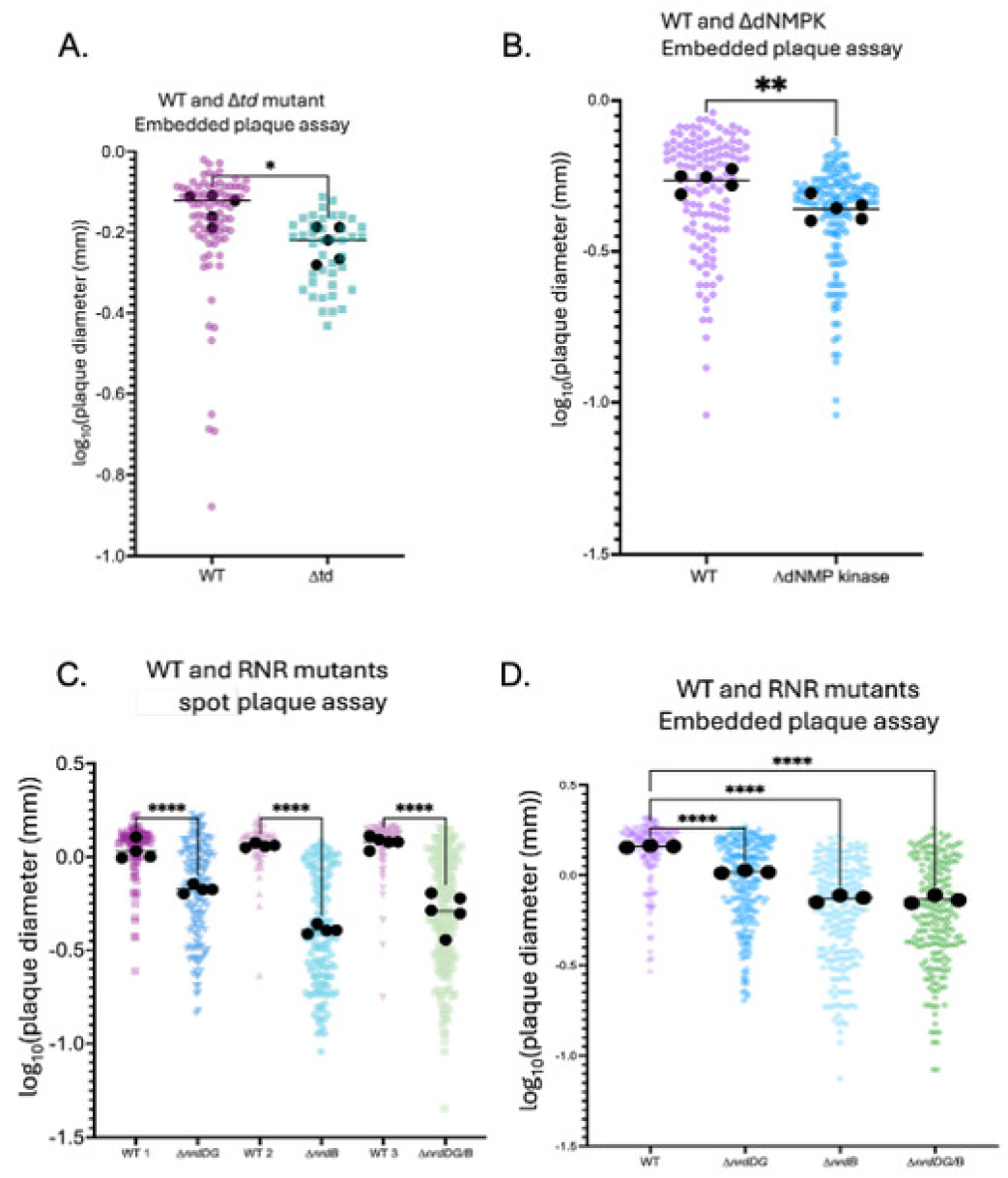
Plaque size assays validate fitness defects identified with Tn-seq. Plaque diameters of deletion mutants of thymidylate synthetase (Δ*td*) **(A)**, gp170 (ΔdNMP Kinase) **(B)**, or ribonucleotide reductase (RNR) genes **(C, D)** were compared with the parent ΔS2-9 strain (referred to as WT in this figure) using either surface (spot) or embedded plaque assays as indicated. Color-filled data points represent individual plaque measurements from 3 to 5 biological replicate experiments, while black-filled circles represent the mean per experiment. Panels C and D include mutants in the anaerobic RNR system (Δ*nrdDG*), the β subunit of the aerobic RNR system (Δ*nrdB*), and an RNR null mutant lacking both systems (Δ*nrdDG/B*). Faded, semi-transparent points represent individual plaque measurements (n = 10 to 50 technical replicates per plate pooled across assays). Large solid symbols represent the mean plaque diameter calculated for each independent biological replicate of embedded plaques (N=3 plates). Normality was confirmed using the Shapiro–Wilk test. We determined statistical significance between groups in panels A and B using a Welch’s t-test on the log10(means), and by a one-way ANOVA performed on the log10( means) for panels C and D (* p < 0.05, ** p < 0.01, *** p < 0.001, **** p < 0.0001, ns = not significant).

Additional nutrient stress hits included gp147 (*helA*; SF1B helicase), gp189 (a putative exonuclease), and gp176 (a putative endoribonuclease) **(Fig. 3B and 3C)**. These genes are encoded in operons with other accessory genes expressed mid-infection to support nucleotide synthesis and replication (37). The putative dNMP kinase activity converts dNMPs to dNDPs, supplying precursors for DNA synthesis. One-step growth curves revealed no difference between ΔdNMPK and wild-type ICP1, despite the plaque-size defect (**Fig. S4**). This repeated trend reflects differences in what each assay captures: plaque size integrates multiple rounds of infection, whereas one-step growth measures a single burst and lysis-timing event, making the former more sensitive to subtle fitness defects.

SF1B helicases such as T4 Dda and ICP1 HelA are implicated in replication, repair, and recombination (38, 39). T4 exonucleases degrade host DNA into oligonucleotides, which can be further processed into mononucleotides, supporting phage nucleotide pools (40). ICP1’s putative exonuclease, Gp189, and putative endodeoxyribonuclease, Gp176, may similarly degrade and scavenge nucleotides from the host or facilitate recombination. In estuary-like media, we also observed depletion of *nrdAB* (gp203-gp204) mutants and *phoH* (gp208) mutants (**Fig. 3C**). PhoH is associated with phosphate acquisition in nutrient-limited environments, and many marine phages encode Pho regulon genes (41). PhoH likely uses ATPase activity to support phosphate uptake, bolstering nucleotide pools under phosphate limitation (31). Some mutants that were depleted after passaging through stationary phase and the estuary-like condition were in hypothetical genes, including gp175, gp024, gp061, gp112 (containing a putative DNA-binding domain), gp119, gp036, and gp037. Interestingly, all but gp036 and gp037 are part of ICP1’s core genome (18). Together, these results show that ICP1 uses auxiliary metabolic genes to synthesize and scavenge nucleotide precursors during nutrient stress when host resources are limited, supporting the hypothesis that ICP1 encodes conditionally important metabolic functions tailored to its ecological niche.

## DISCUSSION

Functional genomics approaches have advanced bacterial genetics, yet tools for studying virulent phages have been limited until recently. Here, we establish a Tn-seq method for the virulent phage ICP1 that enables genome-wide assessment of gene essentiality and conditional fitness during infection of *V. cholerae*. Using an mTn*10* derivative carrying a conditionally essential gene, we generated a nearly saturated mutant library and quantitatively linked phage gene disruption to infection success across diverse growth conditions. The insertion density achieved here allowed us to distinguish a core essential gene set from a large dispensable gene set, revealing that most ICP1 genes are not strictly required for infection under standard laboratory conditions. The essential genes identified that encode proteins of unknown function provide many future research opportunities into ICP1 biology. Because our inducible mTn*10* transposition system works in many other Gram-negative bacteria, this approach may be adaptable to other phage–host systems, provided that a suitable conditionally essential phage gene is available for placing on the mTn*10*.

Our data support the long-standing hypothesis, first proposed for large dsDNA phages such as T4, that accessory genes provide fitness advantages in specific infection contexts (3). Consistent with this hypothesis, Tn-seq and downstream validation revealed that many non-essential ICP1 genes contribute to infection efficiency under nutrient stress or in overcoming hosts that encode specific anti-phage defenses. These findings emphasize that infection ecology shapes phages and that a large fraction of the ICP1 genome may be conditionally useful. Indeed, recent Tn-seq studies in other phages have shown conditionally essential genes needed to infect environmental strains of host bacteria (8–10).

One class of conditionally important genes identified in this study encodes nucleotide metabolism enzymes. ICP1 encodes two RNR systems — the aerobic NrdAB and the anaerobic NrdDG — both of which were classified as non-essential in the input library but showed strong, context-specific fitness defects under nutrient limitation. These two systems behaved similarly, but not identically: Δ*nrdDG* showed a pronounced fitness defect specifically in the embedded plaque assay, which imposes an anoxic gradient, while ΔnrdB showed defects in both embedded and surface (spot) plaque assays. RNR activity is tightly regulated by the availability of cofactors required to generate a free radical for catalysis. NrdAB requires oxygen (32), whereas NrdDG requires S-adenosylmethionine (SAM) and reduced flavodoxin to generate an essential glycyl radical in the NrdD catalytic subunit (33, 34). These separate regulatory mechanisms highlight the advantages of distinct RNR classes and suggest that ICP1 has retained both systems to maintain nucleotide pool balance across a range of oxygen concentrations that it may encounter during infection of *V. cholerae* in different host and environmental niches. These data align with the ecological settings where *V. cholerae* encounters ICP1 predation. The human small intestine represents a rich growth environment for *V. cholerae*. However, upon dissemination in secretory diarrhea to nutrient-poor aquatic environments, the intracellular free dNTP pools collapse as central carbon metabolism slows down (35). Under such growth states and environments, host nucleotide pools are limiting, and phage-encoded auxiliary metabolic pathways may supplement precursor availability to sustain genome replication and maximize burst size.

Many of the dispensable genes with annotated functions in auxiliary metabolism are conserved across sequenced ICP1 isolates, underscoring their importance for phage fitness (18). Indeed, several of these auxiliary metabolic gene mutants were depleted even in the input libraries used to define genome-wide essentiality, which were themselves generated under nutrient-rich laboratory conditions. This suggests these genes confer a general fitness benefit even without nutrient stress.

An important strength of Tn-seq is its ability to identify phage counter-defense genes through direct fitness measurements. Our transposon library construction required ICP1’s CRISPR-Cas activity; accordingly, we showed that all six *cas* genes were conditionally essential. Moreover, using a PLE2 protospacer mutant host in which CRISPR-Cas targeting of PLE2 is abrogated, we demonstrate that Tn-seq robustly detects *adi* as a conditionally important gene specifically required for PLE2 antagonism when CRISPR-Cas-mediated restriction is no longer available. Although the precise biochemical mechanism of Adi remains uncharacterized, it is thought to act by exploiting the PLE2 large serine recombinase to destroy the PLE2 genome (23). This proof-of-principle result confirms that pooled fitness assays can readily capture strong, host-specific counter-defense phenotypes. These results underscore the utility of Tn-seq for identifying phage genes that antagonize known or unknown bacterial immune systems. Since many bacterial immune systems are carried on mobile genetic elements and vary between strains, this approach provides a scalable strategy for uncovering phage adaptations that determine host range and infection success.

While Tn-seq provided a powerful framework to study the importance of ICP1 genes, several limitations became apparent. First, polar effects from promoter-driven transcription within the transposon may alter downstream gene expression, increasing it when in the same orientation and decreasing it when in the antisense orientation. The constitutive promoter embedded in the mTn*10* likely drove the observed directional bias in insertions and may have made antisense insertions deleterious, independent of the disrupted gene’s function. Alternative transposon designs incorporating transcriptional terminators or weaker promoters may mitigate polar effects but could also reduce library complexity, representing a trade-off to consider in future iterations of this method. Second, the pooled, competitive nature of the Tn-seq method yields false positives and noise from population bottlenecks during phage passaging, including the stochastic loss of low-abundance mutants. This distinction matters for interpreting any Tn-seq-based phage screen: the assay is well suited to distinguishing essential, disadvantageous, and neutral mutants at a genome-wide scale, but defects without a strong phenotype should be treated as hypothesis-generating and require orthogonal validation.

When comparing phage Tn-seq with alternative phage functional genomics approaches, several trade-offs are worth noting. Other methods, including CRISPRi-ART, which uses Cas13a to target phage transcripts, and antisense oligonucleotide approaches, offer complementary advantages, particularly for probing essential genes or phages with unusual biology, such as jumbo phages protected by a nucleus-like shell (6, 8, 36). However, these approaches typically require extensive engineering for each new target and are less suited to generating dense, genome-wide fitness landscapes in a single experiment. In contrast, phage Tn-seq provides a high-throughput, relatively unbiased overview of gene importance across the genome.

By revealing how accessory genes shape infection success across environments and host genotypes, this work highlights the dynamic, context-dependent nature of the ICP1 genome and its adaptation to selective pressures imposed by bacterial defenses and nutrient stress. These findings have implications for understanding phage evolution in natural ecosystems and for the rational selection or engineering of phages for therapeutic use. As interest in phage therapy grows, identifying genes that enhance infection robustness, broaden host range, or counter specific bacterial immune systems will be increasingly important. Tn-seq offers a scalable strategy for identifying these determinants at genome scale, while underscoring that arrayed validation remains essential for resolving the precise mechanisms underlying context-dependent phage fitness.

## ACKNOWLEDGMENTS

This work was supported by the National Institutes of Health grants GM139772 (E.A.T.), AI055058 (A.C.) and AI147658 (A.C.).

EAT - Author Contributions: Formal analysis, Investigation, Methodology, Validation, Writing – original draft, and Conceptualization

HBL - Author Contributions: Validation

ACCZ - Author Contributions: Formal analysis, Investigation, and Writing – review and editing

DWL - Author Contributions: Conceptualization, Formal analysis, Investigation, Methodology, Supervision, and Writing – review and editing.

AC - Author Contributions: Conceptualization, Funding acquisition, Project administration, Resources, Supervision, and Writing – review and editing.

